# Exploiting Embryonic Niche Conditions to Grow Wilms Tumor Blastema in Culture

**DOI:** 10.1101/2022.11.07.515473

**Authors:** Heather M. Wojcik, Harold N. Lovvorn, Melinda Hollingshead, Janene Pierce, Howard Stotler, Andrew J. Murphy, Suzanne Borgel, Hannah M. Phelps, Hernan Correa, Alan O. Perantoni

## Abstract

Wilms Tumor (WT), or nephroblastoma, is the most common pediatric kidney cancer. Most WTs display a “favorable” triphasic histology, in which the tumor is comprised of blastemal, stromal, and epithelial cell types. Blastemal predominance after neoadjuvant chemotherapy or diffuse anaplasia (“unfavorable” histology; 5-8%) portend a worse prognosis. Blastema likely provide the putative cancer stem cells (CSCs), which retain molecular and histologic features characteristic of nephron progenitor cells (NPCs), within WTs. NPCs arise in the metanephric mesenchyme (MM) and populate the cap mesenchyme (CM) in the developing kidney. WT blastemal cells, like NPCs, similarly express markers, SIX2 and CITED1. Tumor xenotransplantation is currently the only dependable method to propagate tumor tissue for research or therapeutic screening, since efforts to culture tumors *in vitro* as monolayers have invariably failed. Therefore, a critical need exists to propagate WT stem cells rapidly and efficiently for high-throughput, real-time drug screening. Previously, our lab developed niche conditions that support the propagation of murine NPCs in culture. By applying similar conditions to WTs, we have successfully expanded and passaged WT cells from five distinct untreated patient tumors and maintained key NPC “stemness” markers, SIX2, NCAM and YAP1, and CSC marker ALDH1. These findings suggest that our culture conditions sustain the WT blastemal population, as previously shown for normal NPCs. As a result, we have developed new WT cell lines and a multi-passage *in vitro* model for studying the blastemal lineage/CSCs in WTs. Furthermore, this system supports growth of heterogeneous WT cells, upon which potential drug therapies could be tested for efficacy and resistance.

## BACKGROUND

Wilms Tumor (WT), or nephroblastoma, affects about 1 in 10,000 children, and represents 7-8% of all cancers under age 15 (1). Most WT cases display a “favorable” triphasic histology (FHWT), in which the tumor is comprised of blastemal, stromal, and epithelial compartments, though one cell type may predominate over the others in individual tumors (1,2). In developed countries, overall survival from WT exceeds 90% at 5 years. Relapse affects ∼15% of FHWT patients, among whom roughly half will die, invoking the need for a better molecular understanding of treatment resistance (3). Adhering to Children’s Oncology Group (COG) cooperative trials, therapeutic intensity is risk assigned according to either molecular alterations within FHWT, specifically, loss of heterozygosity (LOH) at both 1p and 16q and at 11p15 or the presence of “unfavorable” histology. Five to eight percent of WT cases display diffuse anaplasia, i.e., 3-fold larger cells with hyperchromatic nuclei and multipolar mitoses, that characterizes “unfavorable” histology (DAWT). This histologic variant alone portends a greater risk for metastasis, relapse, and overall worse outcomes, accounting for half of all WT-related deaths (1,4). Additionally, for children receiving neoadjuvant therapy for FHWT on International Society of Pediatric Oncology (SIOP) protocols, blastemal predominance has been associated with treatment resistance. Further, survivors of WT may display subclinical signs of renal insufficiency and/or treatment-related morbidities later in life. Moreover, survivors 30 to 50 years from initial diagnoses are at increased risk for premature mortality (5,6). Taken together, these factors suggest that current treatment algorithms for WT, although mostly successful, are still not entirely effective and increase the likelihood of secondary health consequences. Thus, there persists a continued need to optimize and personalize initial chemotherapies, tailored to individual WT genetic profiles, and to develop salvage therapies when resistance emerges. To accomplish these goals, however, will require the development of tumor- and cell-specific drug-screening methods for the rapid selection of effective therapeutic agents on an individualized basis, since WT treatment according to COG protocols begins within two weeks of tumor resection, i.e. “real time”. Moreover, a need remains to generate WT cell lines exhibiting the more common forms of favorable histology and various risk-specific mutations, which together associate with the pathologic basis of the disease for research purposes. To this end, establishing WT cell cultures that maintain the more primitive blastemal compartment from primary specimens not exposed to neoadjuvant chemotherapy and upon which initial therapeutics can be tested has proven to be difficult and rarely successful. Whereas some success has been achieved in culturing the epithelial and stromal compartments of WT, effective culture conditions for the blastemal compartment of both FHWT and DAWT have been elusive, and few cell lines exist representing blastemal populations (2,7-10). Typically derived from pre-treated specimens, the currently published cell lines do not display histology representative of the most common triphasic FHWT and either appear anaplastic, harbor TP53 mutations, or exhibit other rare chromosomal abnormalities/mutations that confound general treatment paradigms. Cell lines categorized putatively as WT include MZ128 (truncating mutation exon 2), WiT49 (TP53 mutation, epithelial-like lung metastasis), WT-Pe.1 (chromosomal abnormalities), 17.94 (anaplastic), and PSU-SK-1 (WT1 mutation, fibroblastic) (11-16). These cell lines may be suitable for the study of rare mutations and subtypes but have proven of limited value in the development of therapeutic strategies against the most common forms of FHWT. Furthermore, several published cell lines initially classified as WT have been subsequently reclassified as other tumor types with the emergence of more sensitive molecular diagnostic markers, including G401 (rhabdoid tumor cells), SK-NEP1 (Ewing sarcoma), and WCCS-1 (clear cell sarcoma of the kidney), and are thus unsuitable for the study of WT (17-19). Xenotransplantation of patient-derived WTs has been the only dependable method to propagate tumor tissue for research or therapeutic screening (20,21). However, even xenotransplantation only supports the growth of a fraction of WTs, and, given the relatively slow growth rate of WT implants, is not applicable to “real-time” clinical testing. Therefore, better methods are essential to isolate and propagate WT CSCs/blastemal cells which are amenable to high-throughput, “real-time” drug screening for individualized therapeutic assignment.

The key to the successful *in vitro* culture of WTs without the need for xenotransplantation may lie in the similarity of tumor cells to the normal renal progenitors from which they are derived. WT blastemal cells resemble NPCs of the developing kidney, which arise in the MM and populate the CM. Blastemal cells are also thought to comprise the CSCs of WTs and purportedly marked by ALDH1 and NCAM expressions (21,22). Furthermore, WT blastemal cells also express SIX2, PAX2, and CITED1, which are markers of the self-renewing NPC population (23-29). Due to the histological and molecular similarities between WT blastemal cells and NPCs, it is reasonable to hypothesize that cell culture media capable of stabilizing and propagating NPCs may also support the retention and growth of WT cells in culture.

Previously, our lab developed niche conditions that support the propagation of murine NPCs in culture and demonstrated their dependence on several specific factors for survival and proliferation (30,31). By applying a similar catalog of factors appropriate for NPC cultures to primary WT samples, we have successfully expanded and passaged WT cells which maintain key NPC “stemness” markers, such as SIX2, NCAM, YAP1, and FXR1, as well as the putative WT CSC marker ALDH1. Additionally, cultured cells from one of the WTs successfully grew as a xenotransplant, which could be passaged *in vivo* and which maintained its original histological and molecular profile. Taken together, these data suggest that our culture conditions retain the blastemal/CSC population, as was also observed with our phenotypically and functionally normal cultured NPCs. Thus, using these conditions, we have developed previously unavailable FHWT cell lines and a multi-passage *in vitro* model for studying the blastemal lineage/CSCs in WTs. In light of the rapid and expansive growth of WTs in our cultures, this cell culture approach may finally permit screening for potential drug therapies in “real-time”.

## MATERIALS AND METHODS

### Collection of Primary Wilms Tumor Tissues

The Vanderbilt Institutional Review Board approved all aspects of clinical studies (IRB #’s 100734 and 020888). As published previously (26,28,29), fresh discarded Wilms tumor (VUWT) tissues from patients prior to chemotherapy were provided for processing (laboratory of HNL) and collected in chilled DMEM/F-12 media (Dulbecco’s Modified Eagle Medium; Millipore-Sigma). Tumor cells were either isolated on the day of collection or the next day (stored in media at 4°C until isolation). Briefly, under a sterile hood with laminar flow, whole tumor tissue was transferred to a 10cm cell culture dish containing 20mL Trypsin-LE (Gibco, catalog#12604013) to disperse enzymatically individual WT cells for isolation. Using sterile scissors and conditions, the tissue was minced into small pieces, and the dish with these contents was placed in a 37°C incubator for 30 minutes (32). DMEM/F-12 media was added to the dish, and the solution was filtered through sterile gauze into a 50-mL conical tube to remove non-dispersed tissue debris. The cell suspension was spun down (300xg), and the medium was removed. Cells were washed with DMEM/F-12 media, and the resulting cell pellet was resuspended in ACK Lysing Buffer (KD Medical, catalog# RGC3015) to eliminate red blood cells. The resulting solution was centrifuged at 500xg immediately for 5 minutes. The solution was removed, and the resulting cell pellet was washed with DMEM/F-12. After washing, the cell pellet was resuspended in DMEM containing 10% dimethyl sulfoxide (DMSO) freezing media. Dissociated tissue aliquots were kept frozen at 2 × 10^6^ viable cells/ml/vial in liquid nitrogen until shipment to the National Cancer Institute (author AOP; NCI) and introduction into culture.

### Wilms tumor cell culture

After shipment to the NCI, dissociated tissue aliquots were kept frozen in liquid nitrogen until introduced into culture. For this step, approximately one million cells were seeded onto a 100-mm fibronectin-coated tissue culture dish (Corning BioCoat). Tumor cells were cultured in media similar to that previously reported for the maintenance of murine nephron progenitor cells (NPCs) in tissue culture but modified to include TGF-beta pathway inhibitors used by others (33,34). As we reported previously (30,31), the NPC sustaining media (henceforth referred to as WT media) consisted of a basal medium of DMEM:F12 supplemented with glutamine (GlutaMAX), MEM nonessential amino acids, and Pen/Strep. Several factors which are required for NPC survival and propagation were then added to the basal media, including FGF2, LIF, Y27632, BMP7, CHIR99021, heparin, A83-01, LDN-193189, and fetal bovine serum (FBS). Table 2 details the concentrations and sources of these various reagents. Of note, whereas FBS was not required for murine NPCs, it was essential to maintain these human WT cells but at a relatively low concentration (1% at low density or 2% at high density). GlutaMAX, MNEAA, Pen/Strep, and heparin were diluted in the 500-mL medium container and stored at 4° C. The other factors were added just before application to ensure stability.

### Cell passaging and collection

Cells were grown as both high-density monolayer cultures on 100-mm fibronectin-coated dishes and as organoids in Costar round bottom low adherence 96-well plates or 60-mm dishes incubated at 37° C in 7% CO2. High density semi-monolayer cultures were seeded with 2 million cells per 100-mm fibronectin-coated dish, whereas organoids were grown in either low adherent 96-well plates seeded with 20,000 cells per well or 500,000 cells per low adherent 60-mm dish. Cells in 96-well plates were spun at 50xg to ensure coalescence. Each tumor was cultured in triplicate to allow for statistical analysis; although for some experiments, all three cultures were pooled together. Cells were fed as needed, passaged, and collected at confluence using Accumax:Trypsin (.05%) at 1:1 to dissociate cells. WT cells were collected for flow cytometry and immunoblotting with every passage, from primary tumor samples through passage 5.

### Flow cytometric analyses

WT passages were collected from culture, processed, and analyzed in a Sony SH800 cell sorter. Data from flow cytometry was exported and subsequently analyzed using FlowJo V10 software. To prevent cells from clumping together overnight, which would skew analyses of fluorescent intensities and number of cells analyzed, cells were collected, processed, and analyzed on the same day.

SIX2, a protein expressed by both WTs and NPCs, was analyzed by flow cytometry to discern the percentage of WT cells, at any given passage, which retained SIX2 expression/SIX2 positivity. This allowed us to determine if the culture methods sustained SIX2 expression, which is localized primarily to blastemal WT cells. One million cells were collected per triplicate culture and split in half (500,000 cells) to produce control and experimental populations for each triplicate. Experimental and control samples were permeabilized, fixed, and stained (secondary antibody only for control samples) according to Cell Signaling Technology’s flow cytometry protocol (https://www.cellsignal.com/learn-and-support/protocols/protocol-flow). After staining, cells were evaluated in a Sony SH800 Cell Sorter, and data from at least five thousand cells were collected, where control samples were used to establish proper gating (fluorescence intensity) for SIX2-positive cells. To confirm nuclear localization of SIX2, cultured cells seeded on fibronectin-coated glass slides were also viewed by confocal microscopy.

The ALDEFLUOR assay (Stemcell Technologies Inc., Catalog #01700) was performed using a commercially available kit according to manufacturer’s instructions, including the use of diethylaminobenzaldehyde (DEAB)-treated cells as controls. The assay is based upon an enzymatic reaction to stain for ALDH1 presence and activity in cells. The activity can then be analyzed by flow cytometry. Cells from triplicate cultures were pooled and 2 million cells were collected for staining. This assay only works on viable cells, so no fixation is performed. The ALDEFLUOR metabolite is itself fluorescent and does not require a secondary antibody for analysis. Data were collected using the Sony SH800 instrument and analyzed with FlowJo V10 software. Unstained and DEAB-treated controls were used to determine appropriate gating for positive staining in the experimental samples.

### Immunohistochemistry of Clinical and Experimental Wilms Tumor Tissues

To characterize the expression domain of nephron progenitor markers in the parent clinical and daughter experimental WT specimens, we employed our previously described protocol for peroxidase-based immunohistochemistry (24,26,28,35). Briefly, paraffin blocks containing clinical and experimental WT tissues were cut in 5-μm sections, were subjected to heat-induced epitope retrieval in 10 mM citrate buffer, and were incubated in either affinity-purified rabbit anti-SIX2 (1:20 dilution; US Biological Corp), rabbit anti-NCAM (1:100 dilution; Cell Signaling), rabbit anti-CITED1 (1:50 dilution; ThermoFisher Scientific; Catalog # RB-9219; or Lab Vision/Neomarkers), rabbit anti-YAP (Cell Signaling; catalog # 4912) or mouse anti-PCNA (1:100 dilution; Santa Cruz Biotechnology) antibodies overnight at 4°C. Anti-rabbit or anti-mouse labeled HRP polymer was applied to tissue sections at room temperature for 30 minutes, and antigens were visualized with a DAKO Envision Kit (HRP/DAB System; Dako Cytomation).

For FXR1 detection, slides were placed on a Leica Bond Max IHC Stainer. All steps besides dehydration, clearing, and cover slipping were performed on the Bond Max. Slides were de-paraffinized. Heat induced antigen retrieval was performed on the Bond Max using their Epitope Retrieval 2 solution for 20 minutes. Slides were incubated with anti-FXRl (1:250 dilution; Catalog # - HPA018246, Sigma) for 60 minutes. The Bond Polymer Refine detection system was used for visualization. Slides were then dehydrated, cleared, and cover slipped. All immunostaining was evaluated by a Pediatric Pathologist (H.C.) who was blinded to other clinical or experimental data.

### Immunoblotting

Approximately 4 million cells were collected per passage for protein expression analyses for each tumor. Protein was isolated by incubation of cells in 250 µL of 2X sample buffer (2% SDS, 10% glycerol, 60 mM Tris pH 7.5), boiled at 99°C for 10 minutes, and stored at −20° C until analyzed. Samples (30-µg protein) were loaded in 10% reducing agent, 1:4 dilution of loading dye, and sample buffer, and heated for 10 mins at 99°C. Samples were loaded onto a NuPAGE 4-12% Bis-Tris, 1.5 mm gel (ThermoFisher Scientific) and run for 1 hr at 70 V. Protein transfer onto PVDF filters was accomplished at 70 V for 2.5 hrs on ice for two simultaneous transfers. Filters were blocked in Licor Odyssey PBS Blocking Buffer for 1 hr and incubated at 4° C with rocking overnight in a solution of primary antibody in 10 mL Blocking Buffer and 10 µL Tween-20. All primary antibodies were used at a dilution of 1:1000 except for ß-actin, which was used at a dilution of 1:10000. The next day samples were washed in 0.1% PBST and incubated in secondary antibodies at a 1:20000 dilution in 10 mL Blocking buffer and 10 µL Tween-20. Blots were washed in 0.1% PBST again, dried, and imaged on the Licor Odyssey.

### Xenograft procedures

Assessment of tumorigenicity was accomplished by subcutaneous and/or intrarenal injections of VUWT cells into immunodeficient athymic nu/nu NCr and NSG strains bred in-house. For the subcutaneous implants the cells were mixed with an equal volume of Matrigel or Basement Membrane Extract (Trevigen) and injected subcutaneously on the lateral body wall in a total volume not exceeding 200 µL. The cell density routinely injected was 1 × 10^7^ viable tumor cells. Mice were monitored for general well-being and tumor growth for up to 1.5 years. Initial body weights were recorded so any impact on body weight could be determined as needed. In those cases where a subcutaneous tumor was successfully grown, i.e., VUWT50, the tumor was harvested, cut into small fragments (2×2×2 mm) for passage into a new cohort of mice using a tumor implant trocar. This allowed determination as to whether the tumor could be stably passaged *in vivo*. In the case of the intrarenal inoculation, the standard cell density used was 10^6^ cells/50 µL containing 50% Matrigel, and body weights were serially recorded for comparison. For this step, the mice were anesthetized with isoflurane to effect, the kidney was exposed through an incision just caudal to the rib cage and 5-8 mm off midline to expose the kidney and the tumor cells were slowly injected into the renal parenchyma. The incision was treated locally with bupivacaine and closed with wound clips. Buprenorphine was administered subcutaneously for pain management. Since any potential tumor growth would not be visible, the mice were monitored for general well-being, weight loss, and general signs of disease – imaging was not available. At the time of sacrifice the mice were necropsied and assessed for the presence of lesions. Any lesions observed would have been harvested for serial *in vivo* passage and biological assessment. No tumors were found to occur in or around the implanted kidneys.

## RESULTS

Our laboratory at Vanderbilt has collected numerous, fresh embryonal Wilms tumors (n=54 WT) for research purposes. Seventeen of these WT specimens were enzymatically dispersed immediately after collection for cellular isolation and future culture. From this library of dispersed clinical WT specimens, six were subjected to culture in embryonic niche conditions that support propagation of NPCs. With the intent for separate preliminary cell culture experiments, we determined that VUWT30 grew in similar media and factors as used in our early progenitor experiments. We were encouraged with the propagation of VUWT30 cells in those prior experiments, so the current report describes more robust culture conditions that support rapid and sustainable WT cell propagation *in vitro* and then validates those conditions with *in vivo* growth and replication of the original specimen’s histology. Table 1 shows the patient and disease characteristics of five VUWTs that were representative of the most common FHWT and that we selected to culture for proof of concept. For the purposes of these current studies, we focused on two WT specimens. The first of these cell lines, VUWT13 (Table 1; Figure 1), was established from an early stage, blastemal predominant specimen having perilobar nephrogenic rests that also subsequently relapsed, precisely the scenario of greatest concern clinically. Figure 1 shows the representative histology of the WT tissue acquired in our laboratory after appropriate processing and in parallel to the material used for cell dispersion, isolation, and ultimate culture. This whole WT tissue shows the blastemal predominance at low and high magnification with rare epithelial (i.e., primitive tubules) structures (Figure 1A,B). Panels C-H show detection by immunohistochemistry of various markers of embryonic metanephric mesenchyme and WT blastema (see legend). The patterns of marker expression are consistent with those previously reported (26,27,32,36). VUWT13 principally served to test and optimize niche conditions that would support propagation in culture. The second specimen, VUWT50, was established from an epithelial predominant, stage I tumor not having nephrogenic rests. This case showed numerous tubules and differentiating blastema throughout the acquired specimen. As we have demonstrated in prior work (24,26), CITED1 was robustly detected in blastema with an aberrant nuclear localization, which is typically more cytosolic in the normal embryonic context. Further, some primitive tubular structures in VUWT50, seemingly in zones of active differentiation, indeed expressed CITED1 richly and accumulated in nuclei. Its partner marker of embryonic metanephric mesenchyme, SIX2 (26,37), showed a similar expression domain principally in the nucleus of blastema but also in some primitive tubules as well. Also, as in prior studies (36), FXR1 was intensely detected in the perinuclear cytosol of blastema and again in primitive epithelia in those zones mimicking the embryonic kidney and mesenchymal-to-epithelial transition. After dispersion, isolation, culture, and xenografting of this cell line, blastemal predominant, loosely aggregated Wilms tumors developed in immunodeficient mice, supporting this approach (see below).

**Table 1.** Clinical data for Wilms tumor specimens enzymatically dispersed and cultured.*

| Specimen | Gender | Age (months) | Histology | Cellular predominance | Nephrogenic rests | Lymph Nodes | Stage | LOH 1p/16q | Relapse | Vital Status |
| --- | --- | --- | --- | --- | --- | --- | --- | --- | --- | --- |
| VUWT13 | Male | 44 | Favorable | Blastemal | Yes (perilobar) | Negative | 2 | Unknown | Yes | Alive |
| VUWT22 | Female | 49 | Favorable | Blastemal | No | Positive | 4 | Unknown | No | Alive |
| VUWT34 | Female | 38 | Favorable | Triphasic | Yes (perilobar) | Positive | 4 | No | No | Alive |
| **VUWT50** | **Male** | **56** | **Favorable** | **Epithelial** | **No** | **Negative** | **1** | **No** | **No** | **Alive** |
| VUWT54 | Female | 29 | Favorable | Triphasic | No | Negative | 2 | No | No | Alive |

VUWT50 (bold) is index case analyzed and discussed in text
\* No patient in this study received neoadjuvant chemotherapy, and all cultures were derived from primary, upfront resected specimens according to COG guidelines

**Figure 1.**
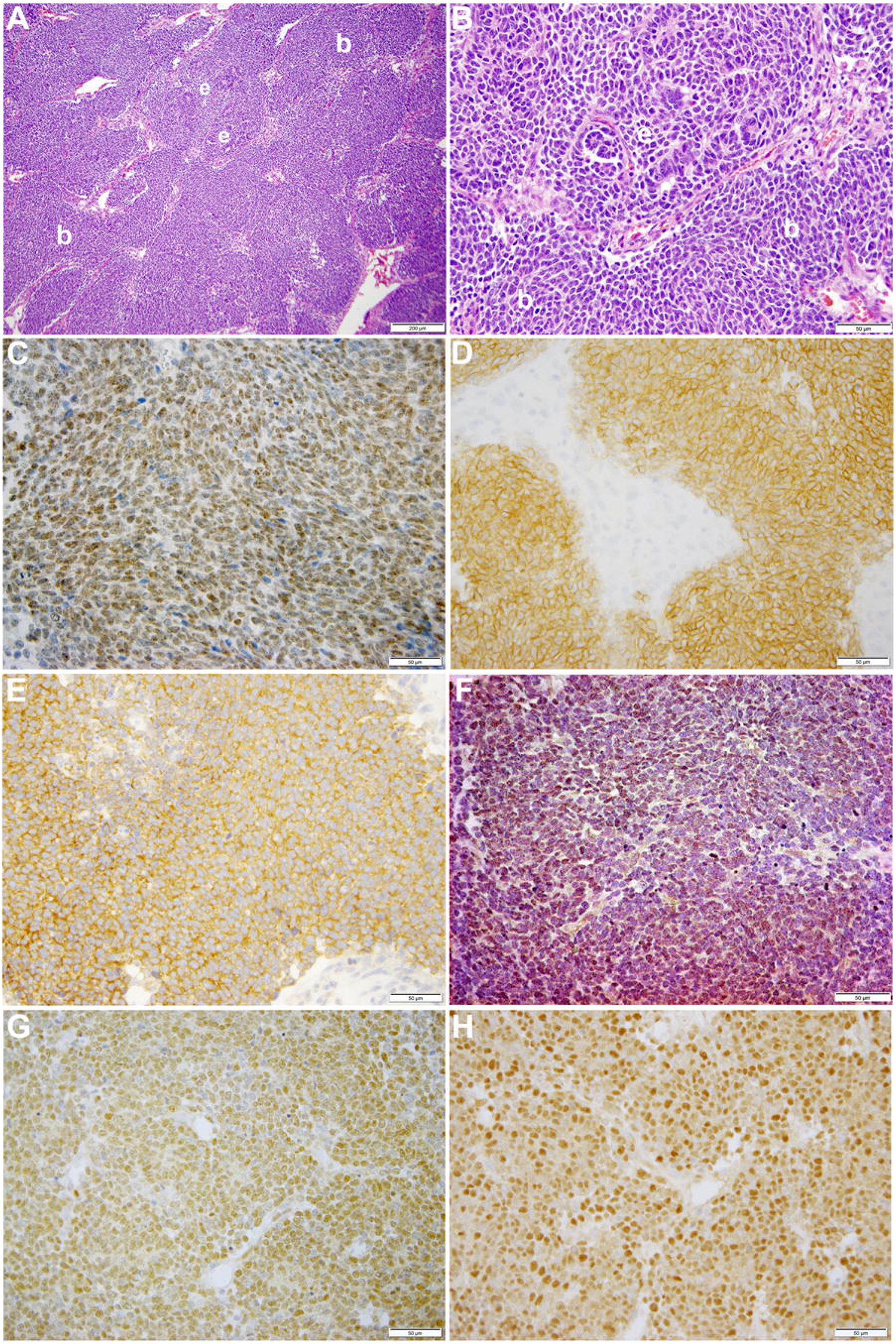
This figure shows representative photomicrographs of VUWT13. *(A, B)* Hematoxylin and eosin stains of primary triphasic, favorable histology Wilms tumor with blastemal predominance. Low- and high-power photomicrographs of same section showing predominant blastema (b) and rare epithelia (e). *(C-H)* These high-power photomicrographs (50 µm bar; 40x) show rich nuclear detection of SIX2 *(C)*, membranous detection of NCAM *(D)*, cytosolic detection of FXR1 *(E)*, nuclear detection of CITED1 *(F)*, nuclear detection of YAP1 *(G)*, and nuclear PCNA *(H)*.

Our previous publications describing the successful cell culture of murine renal progenitors and preliminary work with VUWT30 provide the basis for these current studies (30-32). In those prior studies, we demonstrated the survival and expansion of functional Six2+ nephron progenitors with an ability to differentiate within cell aggregates from both rat and mouse isolated metanephric mesenchymes. Here we assessed the capacity of dissociated WT cells to proliferate and retain stem-like characteristics under similar niche conditions. Each WT culture was initiated from a single vial of liquid nitrogen-stored cells (2 × 10^6^ cells) and plated in the described medium (Table 2) on a 100-mm fibronectin-coated dish. Where possible, aliquots of similarly prepared and thawed cells were applied to flow cytometric analyses. Cells began attaching within an hour and most coated the plate by the following day. The cell growth pattern in cultures of all five tumors was consistent with that observed for VUWT13 (Figure 3A). Cells were small and blastemal-like in appearance, containing nuclei with scant cytoplasm, and retained this morphology despite several passages (Figure 3B). In primary cultures (Figure 3A), larger elongated fibroblasts (presumed stromal cells) that had attached to the matrix were prevalent but appeared less frequently in passaged cells (Figure 3B). The blastemal-like cells preferentially coalesced rather than interact with the substrate as cell density increased. Over time, they formed large 3-dimensional clumps throughout the plate. Given that cell aggregation was essential to the preservation of the renal progenitor phenotype in normal murine tissues, tumor cells were grown to extremely high densities to encourage cell-cell interactions. Due to their rapid doubling times (from 0.35 to 1.0 average doubling/day) though, cells required passaging at least once every 6-7 days, yielding cell numbers ranging from 20-50 million cells/100 mm dish. All five cultured VUWTs were carried in this manner for at least five passages to assess the stability of their progenitor-like phenotype. Sufficient cultured cell numbers were produced for xenotransplantation studies by passage 3 (P3) for all five tumors.

**Table 2.**
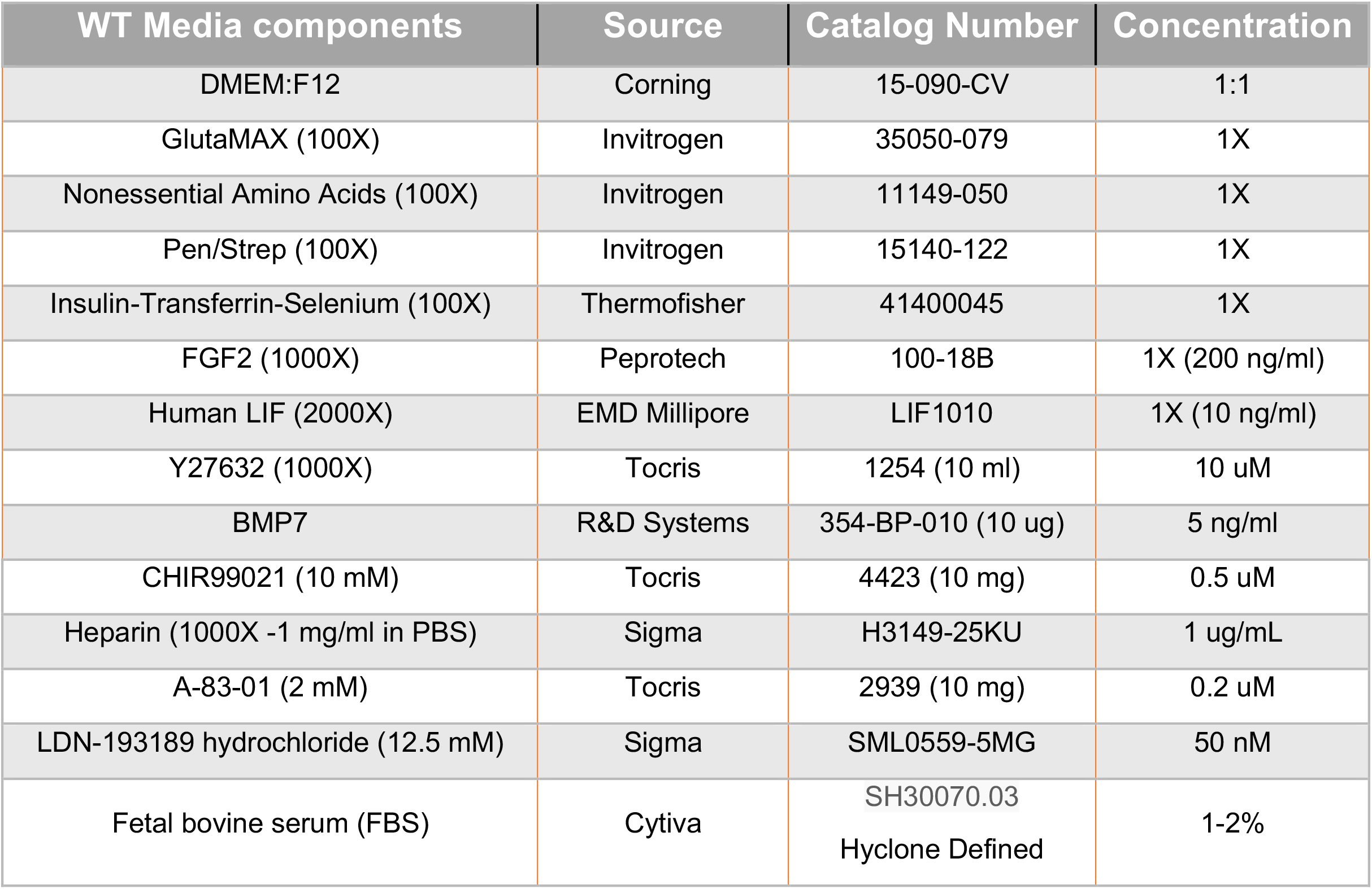
Medium composition and reagent source.

To directly assess the presence and stability of a key progenitor marker Six2, we performed flow cytometry on cells from all passages and for all five tumors. Three independent cultures were generated from Passage 0 (P0) cells and carried separately throughout the study. For VUWT13 and VUWT50, Six2+ cells typically represented more than two thirds of the cells analyzed (Figure 3C) throughout passaging. The actual scatter plots used to generate the composite are shown in Supplementary Figure 1. Flow cytometric assessments of all passages from VUWT22, VUWT34, and VUWT54 yielded similar results with averages of Six+ cells typically representing 60-70% of the cell population (Supplementary Figure 2). These data demonstrate that cultured WT cells retain their progenitor-like phenotype with the expression of Six2 and also that cultures consist of mixed cell populations as expected from tumors with favorable histology. To demonstrate that expression of SIX2 was predominantly nuclear localized as shown in sections from fixed patient tumor tissues (Figure 2), VUWT13 p5 cells were analyzed by immunofluorescence labeling (IF), which confirmed nuclear staining (Figure 3D). Cells were also probed for YAP1 by IF and observed to express this tumor marker in cultured tumor cell nuclei as well (Figure 3E). This observation corresponds with the staining pattern in fixed tumor tissues (Figure 2 and prior report (32) and is consistent with YAP activation in tumor cells (38,39).

**Figure 2.**
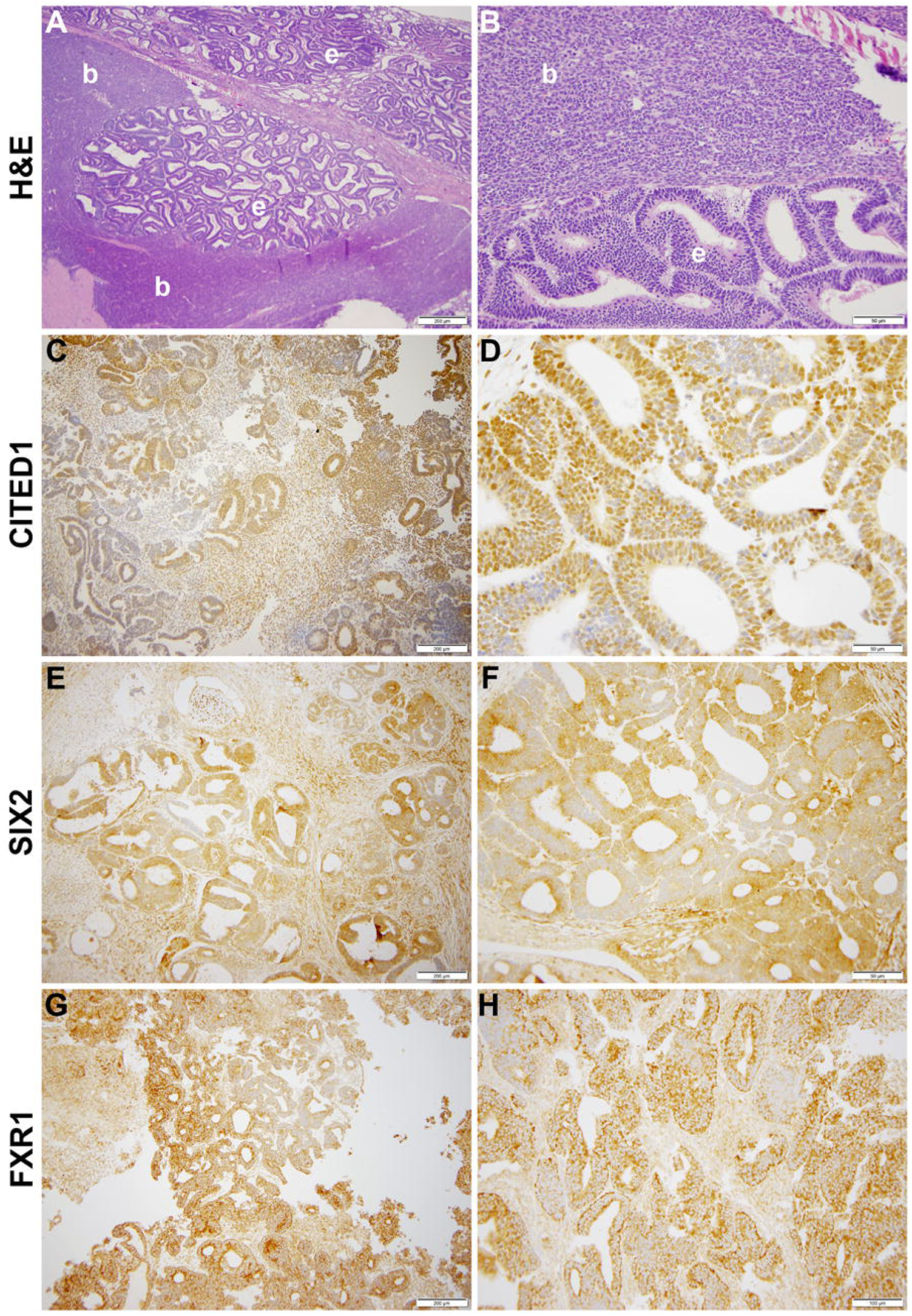
This figure shows representative photomicrographs of VUWT50. *(A, B)* Hematoxylin and eosin stains of primary triphasic, favorable histology Wilms tumor with epithelial predominance and heterologous differentiation. Low- and high-power photomicrographs of same section showing blastema (b) and epithelia (e). *(C, D)* CITED1 detection in nearly all blastema but roughly half of primitive epithelia (*A:* 200 μm bar, 10x; *B:* 50 µm bar, 40x). Note subcellular localization is intense in nuclei but also cytosol. *(E, F)* SIX2 detection in blastema and primitive epithelia (*C:* 200 μm bar, 10x; *D:* 50 µm bar, 40x). Subcellular localization parallels CITED1. *(G, H)* FXR1 detection in blastema and epithelia, with subcellular localization principally along cytosolic perinuclear membrane (*E:* 200 μm bar, 10x; *F:* 100 µm bar, 20x).

**Figure 3.**
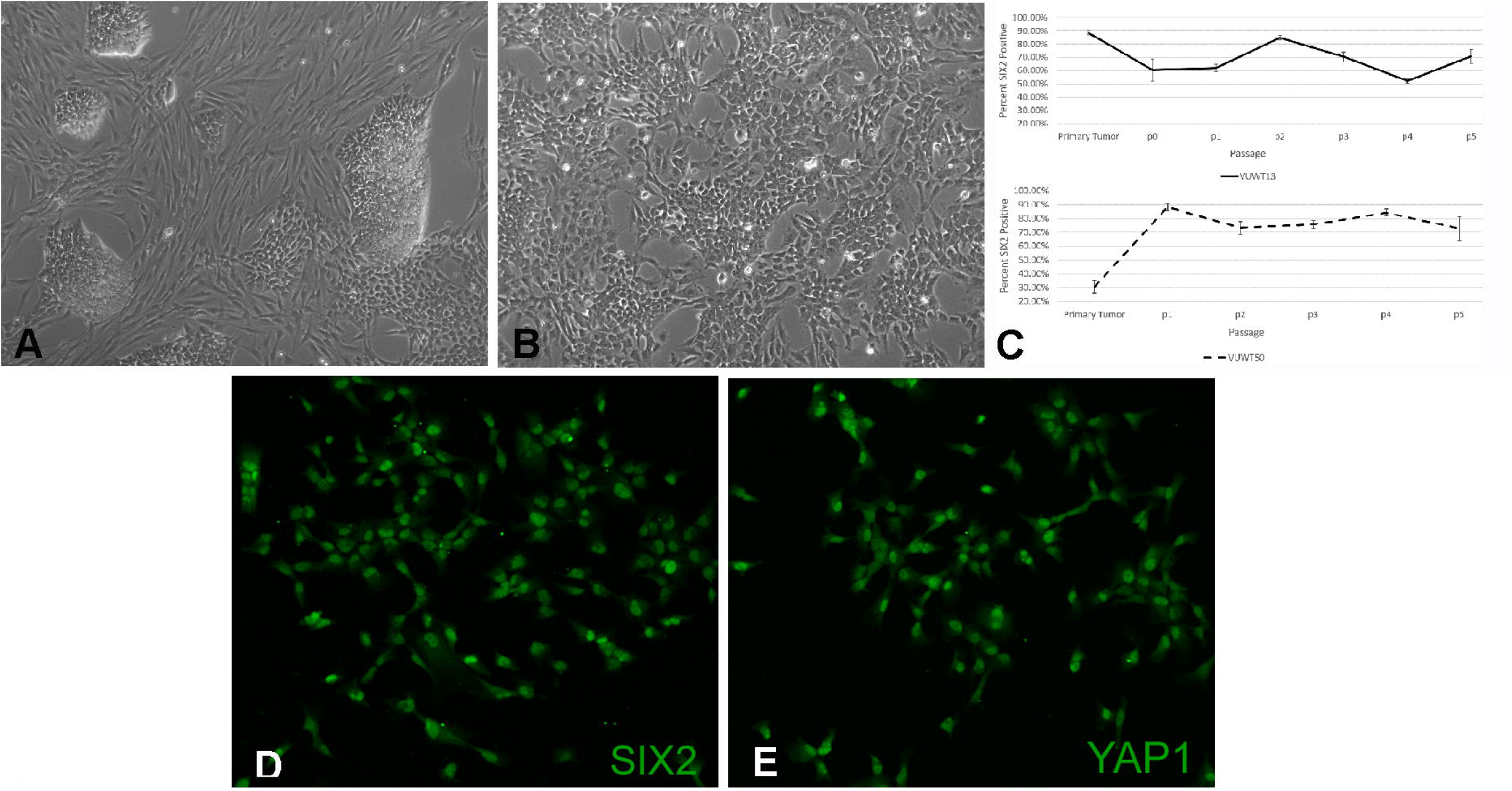
Analysis of cultured VUWT13 or VUWT50 tumor cells. A. VUWT13 cells in culture directly from frozen tumor tissue, showing a mixed population of blastemal and stromal cells. B. VUWT13 cell morphology in passage 5 cultures, showing primarily blastemal cell populations. C. Flow cytometric analyses for Six2 expression in VUWT13 and VUWT50 through passage 5 cultures. D. Immunofluorescence staining for SIX2 expression in p5 cultures of VUWT13 cells is predominantly nuclear localized. E. Immunofluorescence staining for YAP1 expression in p5 cultures of VUWT13 cells shows prominent nuclear localization. F. VUWT13 and VUWT50 cultures contain small but stable populations of ALDH1+ cells as determined by an Aldefluor flow cytometric analysis.

ALDH1 is identified as a potential cancer stem cell marker in a variety of neoplasms, including WT, and high expression or activity is associated with a poor clinical outcome in some cancers (40-44). Recent studies suggest that ALDH1 functions downstream of ß-catenin-dependent Wnt signaling (43,44), which is often constitutively active in WTs as well (45,46). To determine if the current VUWTs also manifest ALDH1 activity, we employed an ALDEFLUOR assay for flow cytometric identification of the ALDH1+ subpopulation. Using this assay, we determined that both VUWT13 and VUWT50 contained significant subpopulations of ALDH1+ cells and that those cells remained relatively stable with passaging (Figure 4). Notably, the majority of cells in the predominantly blastemal tumor VUWT13 were ALDHi+. VUWT34 and VUWT54 contained small but measurable percentages of ALDH1+ cells, which remained relatively stable throughout passaging (Supplementary Figure 3). For VUWT34, the subpopulation initially declined but recovered by P3 and remained stable thereafter. These data further demonstrate that our niche culture conditions maintain a mixed cell population and suggest that the putative CSC has been retained in our cell cultures regardless of passage. These percentages of ALDH1+ cells in the current experiments were consistent with reports from other studies of WT (21,47,48).

**Figure 4.**
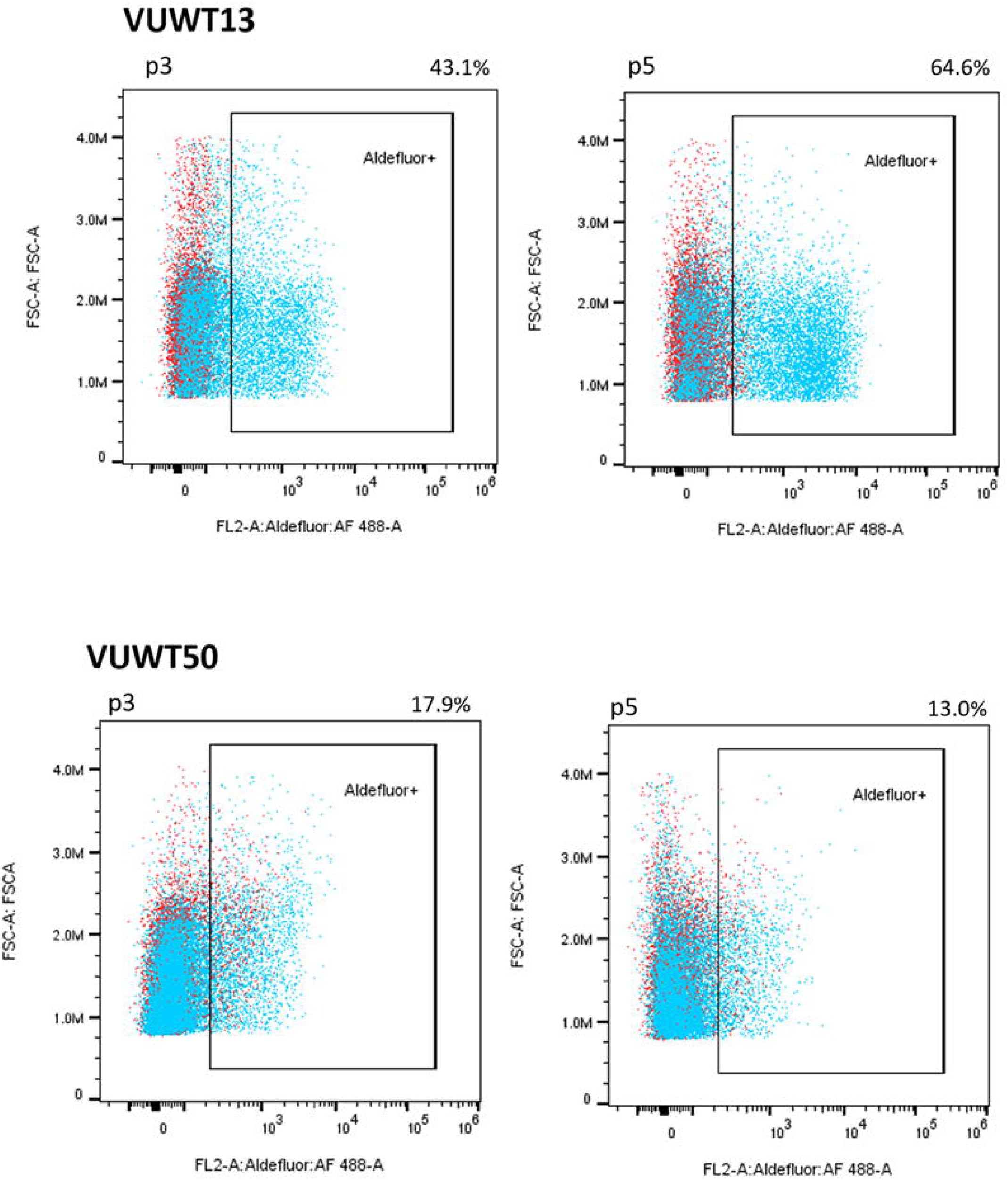
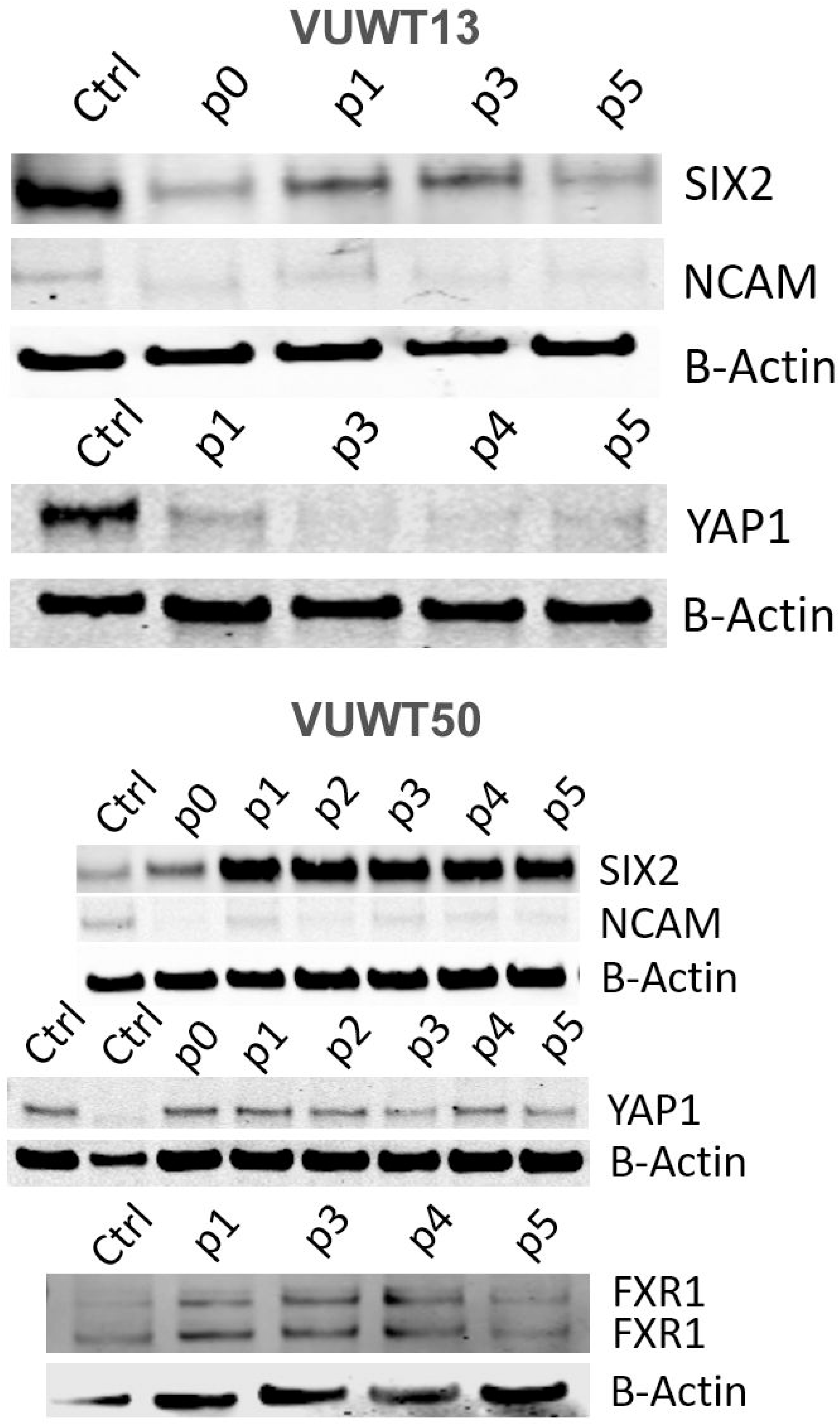
Immunoblot analyses of VUWT13 and VUWT50 for the expressions of common Wilms tumor markers, SIX2, YAP1, NCAM, and FXR1. All markers were demonstrable through multiple culture passages with some variation.

To further assess the presence and stability of expression of various WT markers, we applied immunoblot analyses for SIX2, YAP1, NCAM, and FXR1 (36,49). All four markers were demonstrable in most tumors included in this study (Figure 5 for VUWT13 and VUWT50 and Supplemental Figure 4 for VUWT22, VUWT34, and VUWT54) and in every passage, albeit at varying levels in different tumors. Overall levels of SIX2 and YAP1 tended to be highest in the earlier passages in immunoblots; however, both remained detectable even in later passages. NCAM levels were consistently observed in these studies apart from VUWT34 and VUWT54, where expression was indeterminate at the protein level. FXR1 was not included when VUWT13 was first analyzed; however, it was detected subsequently in spheroids, as shown later.

**Figure 5.**
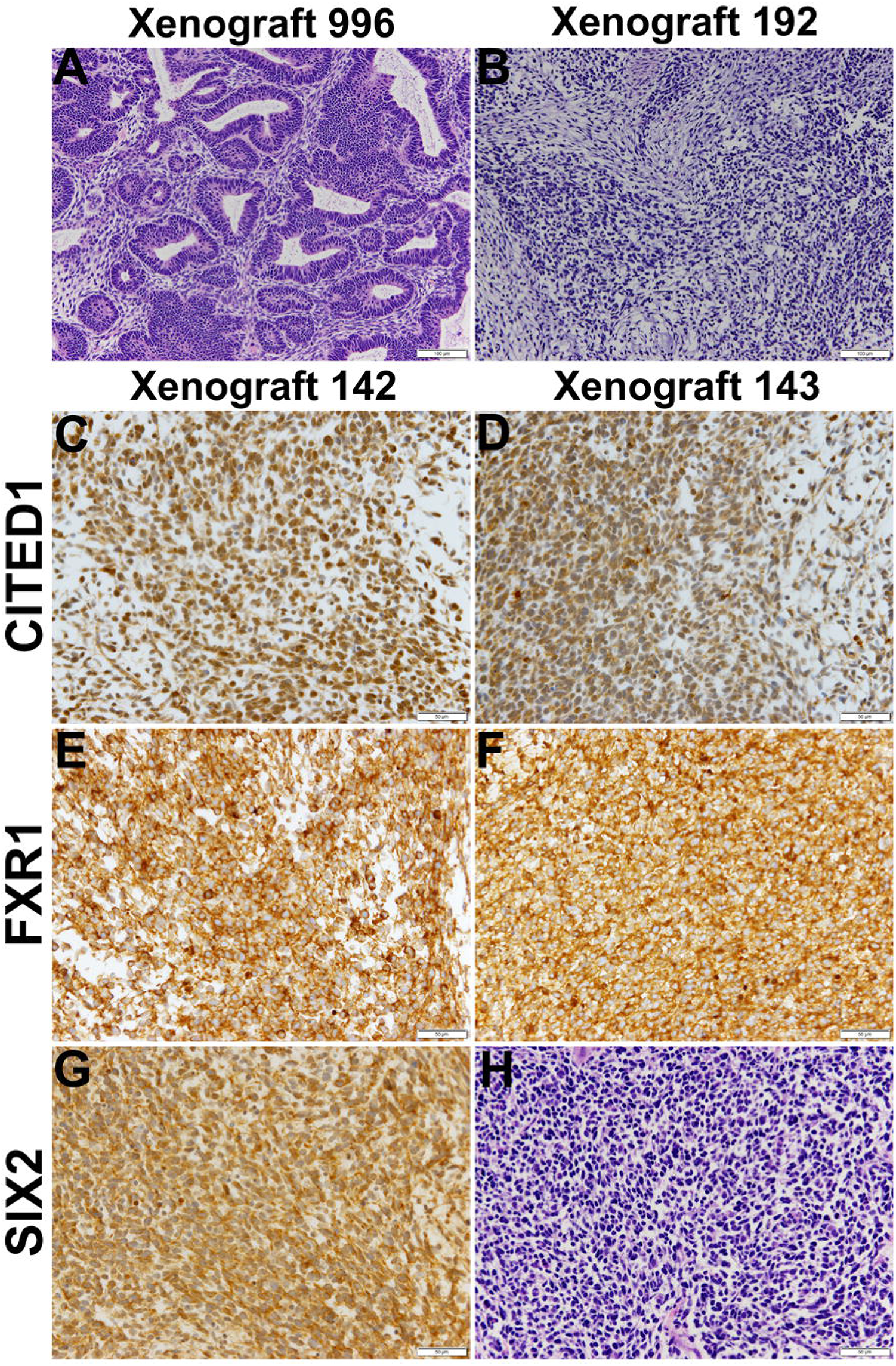
This figure shows representative histology and immunostaining of VUWT50 cells after culture, then engraftment into subcutaneous tissues of nude mice. *(A, B)* Hematoxylin and eosin stains of passage 2 (996) and 3 (192) xenografts originating from VUWT50. Each column for either xenograft represents a 10x (bar is 200 µm) and 40x (bar is 50 µm) photomicrograph. First passage xenografts of previously cultured VUWT50 cells show classic triphasic Wilms tumor identical to primary resected specimen. Xenograft 192 shows greater cellular predominance of blastemal elements that are loosely aggregated and producing a myxoid substance, typical of Wilms tumor blastema. *(C, D)* CITED1 detection in two distinct passage 3 xenografts (142 and 143) from VUWT50 (40x; bar is 50 µm). CITED1 shows retained rich cytosolic and nuclear expression and is restricted to the blastema of each specimen. This robust CITED1 detection shows maintenance of the blastemal element with later xenograft passage. *(E, F)* FXR1 detection is robust and cytosolic, with a unique intensity in the perinuclear location (40x; bar is 50 µm). *(G)* SIX2 shows strong nuclear and cytosolic expression (40x; bar is 50 µm). *(H)* The H&E shows blastemal-predominant histology, which was representative of both xenografts 142 and 143.

To determine their tumorigenic potential, cultured cells, primarily at P3, from each tumor were suspended in soft agarose, containing the same factors applied to monolayer cultures and maintained for up to 6 weeks to allow for colony formation. For colony counts, only intact multicellular aggregates were included. As documented in Table 3,all five tumors formed colonies to varying degrees; however, only VUWT13 and VUWT34 produced substantial colony numbers. These data indicate that passaged cells have maintained an ability for anchorage-independent growth, a property consistent with a transformed phenotype.

**Table 3.** Colony formation in agarose by cultured Wilms tumor cells.

| Wilms tumor | Colony count/50K cells |
| --- | --- |
| VUWT13, p5 | 145 ± 20 |
| VUWT22, p3 | 1 ± 0.3 |
| VUWT34, p3 | 285 ± 20 |
| VUWT50, p3 | 6 ± 3 |
| VUWT54, p3 | 4 ± 0.6 |

Historically, xenotransplantation has proven the only effective method for the maintenance and growth of isolated WTs (35,47). To further assess their tumorigenic potential, P3 cells from all five dispersed VUWTs were applied to xenotransplantation studies in immunocompromised mice (using both athymic nu/nu NCr and NSG strains), as described in Materials and Methods. Following subcutaneous or intrarenal injections, five mice for each tumor preparation and injection site were assessed for tumor growth for up to 1.5 years. Of these, cultured cells from VUWT50 p3 established tumors in 3 of 5 implanted athymic nude mice (nu/nu NCr) and 1 of 5 NSG mice after subcutaneous injection. Tumors occurring in both mouse strains were successfully serially passaged into new hosts through 4 passages, at which time, we discontinued passaging as the tumor was stably preserved as a human tumor. Xenotransplantation passage 2 (xenograft 996) and passage 3 (xenograft 192) tumors upon histologic examination (Figure 6) showed classic features of triphasic FHWTs, consistent with the primary clinical specimen (Figure 1). A Pediatric Pathologist (HC) was blinded to experimental protocol and diagnosed each of these xenograft specimens as WT and as retaining the histologic features of the parent specimen. This Pediatric Pathologist further reviewed all immunostaining for the examined molecular markers of WT blastema. Additional passage 3 specimens (xenografts 142 and 143) showed blastemal-predominant histology and stained robustly for CITED1, a marker of proliferating WT blastema (Figure 6 C,D, and (50)). These latter two xenografts (passage 3), also showed robust expression of FXR1 (Figure 6 E,F) and SIX2 (Figure 6 G,H). Of note, CITED1 was again detected prominently in nuclei from all specimens but also exhibited some cytosolic staining, as previously shown in primary WTs (24). FXR1 was detected in the cytoplasm with a rich perinuclear expression, also as shown previously (36). SIX2 was observed in both the cytosol and nuclei in these xenografts. Both histology and immunohistochemistry of VUWT50 are consistent with the nature of the patient’s original tumor (Figures 1 and 2).

**Figure 6.**
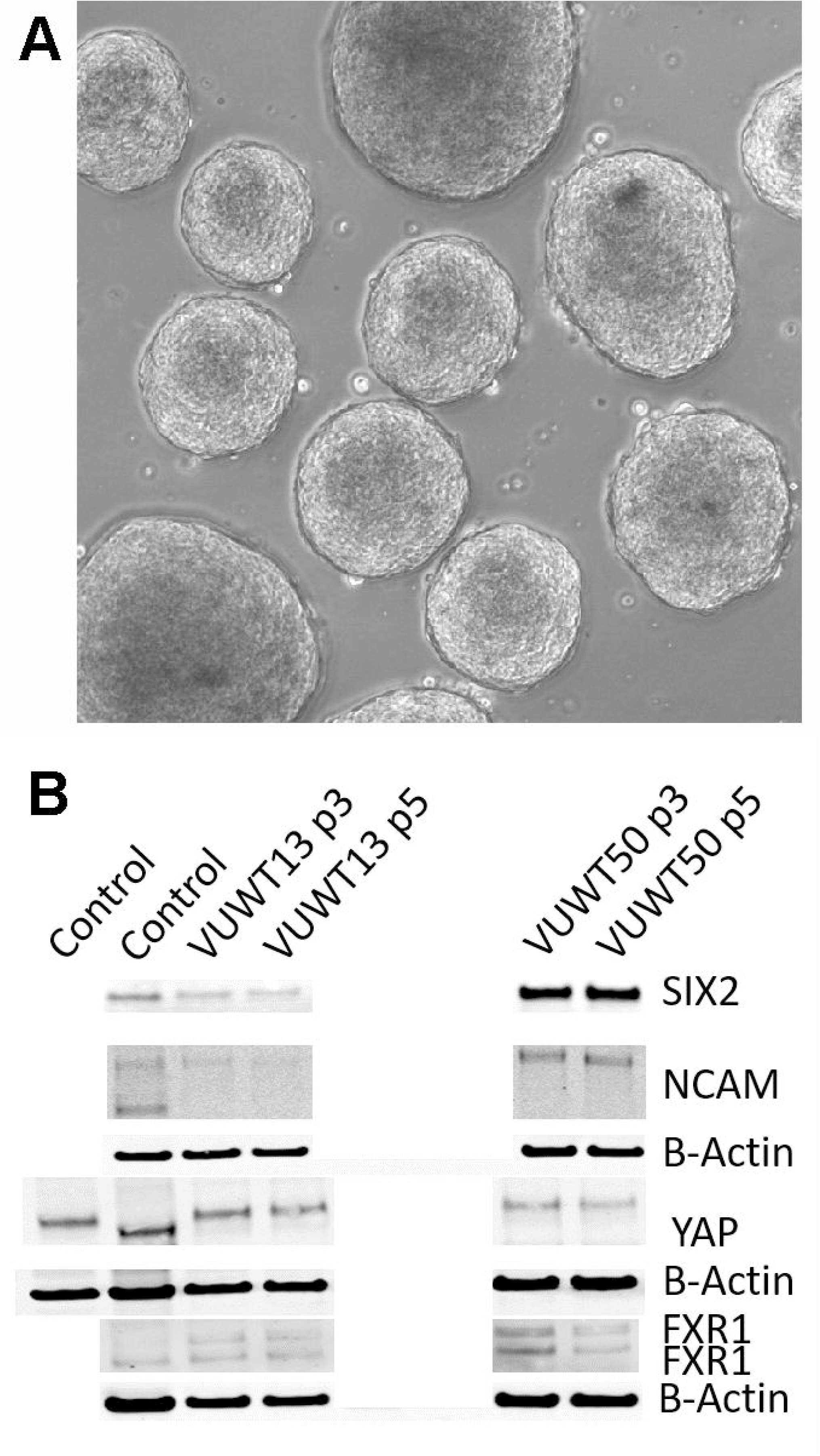
Cultured tumor cells readily form spheroids in the current medium conditions. A. VUWT13 cells form spheroids in wells of non-adherent culture plates and could be maintained with passaging. B. Immunoblot analysis demonstrated the stable retention of expression of Wilms tumor markers in spheroids from VUWT13 and VUWT50 cells that were dissociated and re-aggregated through several cycles. HEK293 or whole mouse embryonic kidney lysates were used as controls.

Recently, considerable interest in tumor cell culture has focused on the ability of tumor cells to form spheroids on low adherence culture plates. Indeed, WTs have also been reported to form spheroids/organoids under such conditions (51,52). To determine if our current culture conditions will also support the formation of similar aggregates, we suspended 20,000 cells in each of several wells of a 96-well low adherence plate. The plate was then spun at 50xg to assist in cell aggregation. When enlarged aggregates began sloughing cells (at about 100,000 cells), spheroids were collected, dissociated, and resuspended for the next passage. This typically required 8-12 days of growth for each passage. VUWT13 spheroids readily formed in wells, and several are shown in Figure 7A. When analyzed for expression of stem-like markers by immunoblot (Figure 7B), VUWT13 and VUWT50 expressed all markers examined (SIX2, YAP1, NCAM, and FXR1) in both P3 and P5 cells, as shown earlier for 2-D cultures from these cells. Similarly, VUWT22, VUWT34, and VUWT54 yielded a pattern of expression for these proteins that reflected our observations for 2-D cultures (Supplemental Figure 5). These observations indicate that our culture conditions support the formation and maintenance of WT cells as spheroids.

**Figure 7.**
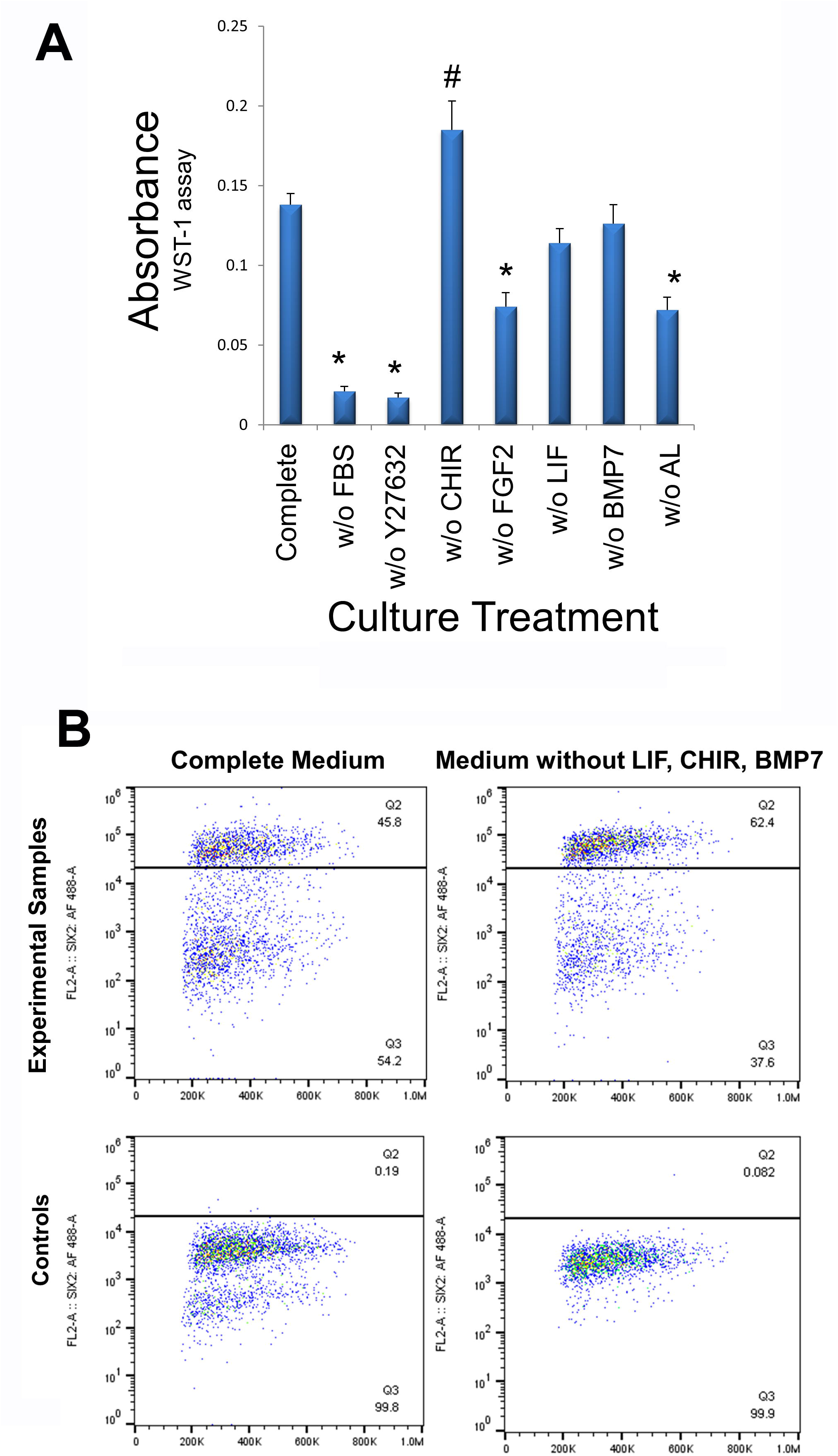
Determination of media components essential for cell growth. A. VUWT13 cells at p5 were cultured in 24-well plates for 4 days with the indicated treatments. Cells were then exposed to the WST-1 reagent for 4 hours and absorbance measured at 440 nm. FBS (1%), ROCK inhibitor Y27632, FGF2, the combination of ALK inhibitors all promote growth in this assay. *Growth is significantly reduced relative to controls (p<0.05); # Growth is significantly increased relative to controls (p<0.05). B. To determine if the tumor phenotype remained stable despite the absence of factors that failed to promote growth, we also quantified the percentage of P10 SIX2+ cells in cultures by flow cytometry, as shown in scatter plots. The percentage of SIX2+ cells did not decrease in cultures that had been maintained in the absence of non-growth-promoting factors despite multiple passages.

Finally, all media components used in cultures of murine NPCs with the addition of 1% FBS and two Alk inhibitors were included in the culture of WT cells in the current study. Since normal progenitors were dependent upon these factors to maintain their progenitor phenotype, we applied the same factors to WT cells as well. However, it may prove advantageous economically to reduce the niche condition complexity by eliminating some of the costly factors, if they play no significant role in the growth of WT cells and maintenance of the stem-like phenotype. Accordingly, we assessed the growth of VUWT13 cells following the deletion of individual media components. In Figure 8A, we observed that the critical factors promoting growth in our media are FGF2, Rho kinase inhibitor Y27632, FBS, and the two Alk inhibitors. The factors without apparent effect on growth include leukemia inhibitory factor (LIF), Wnt agonist CHIR, and BMP7. In fact, the CHIR appeared to inhibit growth. While these data indicate that a less complex medium can support tumor cell growth, they do not inform us on the stability of the tumor phenotype. Accordingly, we also evaluated cells maintained separately over time in our initial complete medium or in medium lacking LIF, CHIR, and BMP7, which showed no significant effect on growth. By flow cytometry of VUWT13 p10 cells, we observed, not only that our original conditions maintained high levels of SIX2+ cells, but also that elimination of CHIR, BMP7, and LIF did not reduce or eliminate the percentage of SIX2+ cells in cultures of VUWT cells (Figure 8B). These findings suggest that the less complex medium may be sufficient to support the rapid growth and maintenance of WT cells, but further studies using primary tumor cells will be required to validate this approach.

## DISCUSSION

In the current study, we successfully culture and grow cells from five distinct WTs, all with favorable histology, and demonstrate that the cultured tumor cells retain a predominantly blastemal-like phenotype, revealing expressions of key genes such as SIX2, YAP1, NCAM, and FXR1 through several passages using flow cytometry or immunoblotting. Furthermore, SIX2 and YAP1 remain nuclear-localized, as observed for normal nephron progenitors, and suggesting that oncogenic YAP1 is constitutively active in these tumors (32,53). Also, ALDH1 activity is detected in significant numbers of tumor cells. Given that ALDH1 expression appears to be downstream of Wnt signaling (43,44), which has also been implicated in WTs, it is possible that it serves only as a marker for Wnt pathway activation. Still, ALDH1 is frequently associated with clinical progression for a variety of human cancers and can therefore serve to identify potential CSCs within these tumors. Importantly, despite the small percentages of ALDH1+ cells within some tumors, those numbers were relatively stable and resolvable due to the ability for analysis by flow cytometry.

As for properties of Wilms tumorigenicity in our cultured cells, we also show that these cells maintain an ability for anchorage-independent growth and, for at least one specimen tested, that cultured cells have the capacity to establish tumors when xenotransplanted, which has never been reported for 2-D cultured WT cells. It may be noteworthy that the two tumors that contained nephrogenic rests, i.e., VUWT13 and VUWT34, yielded by far the largest number of colonies in agar assays. Moreover, the one tumor which grew in xenotranplants, i.e., VUWT50, was derived from a predominately epithelial tumor. Whether the biology of these tumors is relevant to their ability to grow under the conditions imposed upon them by these assays remains a matter of speculation though, due to the small sampling size and requires additional studies.

To date, as indicated in the Introduction, the culture of WTs has been challenging and largely unsuccessful, precluding the possibility of an individualized approach for therapeutic intervention. This current novel study, however, provides evidence to support the possibility for “real-time” clinical testing of WTs, not only for diagnostic purposes, but also for chemotherapeutic screening. A critical factor in such approaches is the stabilization, rapid growth, and retention of the original tumor histologic and molecular pathology. Our innovative approach to exploit embryonic niche conditions successfully addresses these considerations. The principal population of cells preserved in cultures displays phenotypic traits of the nephron progenitor and primary patient tumor tissues. Such ancestral traits may represent the critical driving force for tumor development and retention of stem cell properties. For example, the ALDH1+ population is associated with a negative clinical outcome and can be quantified in our culture system, thus permitting the selective screening for agents that preferentially target the ALDH1+ WT cells. Importantly, the rapid proliferation of cells in culture using our niche conditions yields massive numbers of cells for a variety of simultaneous analyses within the first two weeks following tumor excision. Within that time window, it would theoretically be possible to develop personalized therapies for each individual patient. To put this in perspective, the WT volume at diagnosis reported in one study averaged 570 ml (54). Similar tumor cells display a cell density slightly greater than 1 gm/mL (55), thus the average tumor weighs about 570 grams. Since one gram of tumor tissue is composed of 10^8^ to 10^9^ cells (56), conservatively it should be possible to harvest 2 × 10^10^ cells from that tumor for therapeutic screening. Current high-throughput screening (HTS) platforms utilize 384-well dishes with 10,000 cells/well or nearly 4 × 10^6^ cells/plate (57). In that regard then, the limiting factor in HTS studies would be the cost of screening 5000 plates and not cell numbers. A significant problem, however, has been the inability to stabilize and culture cells from individual patients for the duration of the screening process. Recent successes in culturing spheroids from patients’ tumors has attempted to address this problem. There is indeed a basis for employing spheroids/organoids in screening studies to provide a more realistic 3-dimensional structure, since such structures may limit drug penetration and thus drug efficacy. However, cell growth in spheroids is nearly an order of magnitude slower with doubling times of 1-2 weeks in one set of conditions (51) or with 1:3 splits every 10-14 days (52). Our spheroid growth more closely resembles those earlier reports, i.e., 5-fold increases in cell numbers over 8-12 days for spheroids versus 25-fold increases over 6 days for our 2-D cultures. Under the current culture conditions, a preliminary 2-D screening could be performed during the first week of “real-time” testing with a follow-up in spheroid cultures on a more targeted basis the subsequent week. Having the flexibility to apply both approaches enhances the utility of our conditions. Furthermore, the faster growth rates in 2-D screening are likely to reveal significant endpoint differentials more rapidly as well, further supporting the efficacy for “real-time” testing using our culture conditions. To that end, it is now possible to validate this approach with the niche conditions described in this report.

## ACKNOWLEDGEMENTS

The authors gratefully acknowledge Carrie Bonomi and Kelly Dougherty Leidos Biomedical Research, National Cancer Institute at Frederick, Frederick, MD, USA for excellent technical assistance in the xenotransplant studies. They also acknowledge Sherry Smith, Kelly Boyd, Cindy Lowe, and the Vanderbilt Translational Pathology Shared Resource for technical assistance in performing portions of the immunohistochemistry experiments. This research was supported in part by the Intramural Research Program of the NIH, NCI, Cancer and Developmental Biology Laboratory. The Translational Pathology Shared Resource at Vanderbilt University is supported by NCI/NIH Cancer Center Support Grant P30CA068485. This work was supported in part by NCI grants 4R00CA135695-03 (HNL), 5T32CA106183-06A1 (AJM), and by the Section of Surgical Sciences and the Ingram Cancer Center of the Vanderbilt University Medical Center. Additional funding was obtained from the National Center for Advancing Translational Sciences [CTSA grant number UL1 TR002243 (HNL)]. This project has been funded in part with federal funds from the National Cancer Institute, National Institutes of Health, under Contract No. HHSN261200800001E. The content of this publication does not necessarily reflect the views or policies of the Department of Health and Human Services, nor does mention of trade names, commercial products, or organizations imply endorsement by the U.S. Government. This research was supported in part by the Division of Cancer Treatment and Diagnosis of the National Cancer Institute. NCI-Frederick is accredited by AAALACi and follows the Public Health Service Policy on the Care and Use of Laboratory Animals. All animals used in this research project were cared for and used humanely according to the following policies: The U.S. Public Health Service Policy on Humane Care and Use of Animals (1996); the Guide for the Care and Use of Laboratory Animals Eighth Edition; and the U.S. Government Principles for Utilization and Care of Vertebrate Animals Used in Testing, Research, and Training (1985).

## Figure Captions

**Supplementary Figure 1**. Scatter plots of flow cytometric quantification for SIX2+ VUWT13 and VUWT50 cultured and passaged cells. Controls included only secondary antibody for proper gating.

**Supplementary Figure 2**. A composite graph of flow cytometric analyses of SIX2 expression in all passages of cultured VUWT22, VUWT34, and VUWT54 cells.

**Supplementary Figure 3**. Analysis of ALDH1+ cells for cultured passages of VUWT34 and VUWT54 cells using flow cytometry.

**Supplementary Figure 4**. Immunoblot analyses for Wilms tumor markers in cultured passages of VUWT22, VUWT34, and VUWT54 cells.

**Supplementary Figure 5**. Immunoblot analyses for Wilms tumor markers in spheroids from various cultured passages of; A - VUWT22, B - VUWT34, and C - VUWT54 cells. VUWT13 and VUWT50 were taken from these blots and included in Figure 6. HEK293 and embryonic mouse kidney were used as controls.

## Notes

### Competing Interest Statement

The authors have declared no competing interest.

