## Supplementary figures and images for "Exploiting Embryonic Niche Conditions to Grow Wilms Tumor Blastema in Culture"

### Supplementary Figure 1

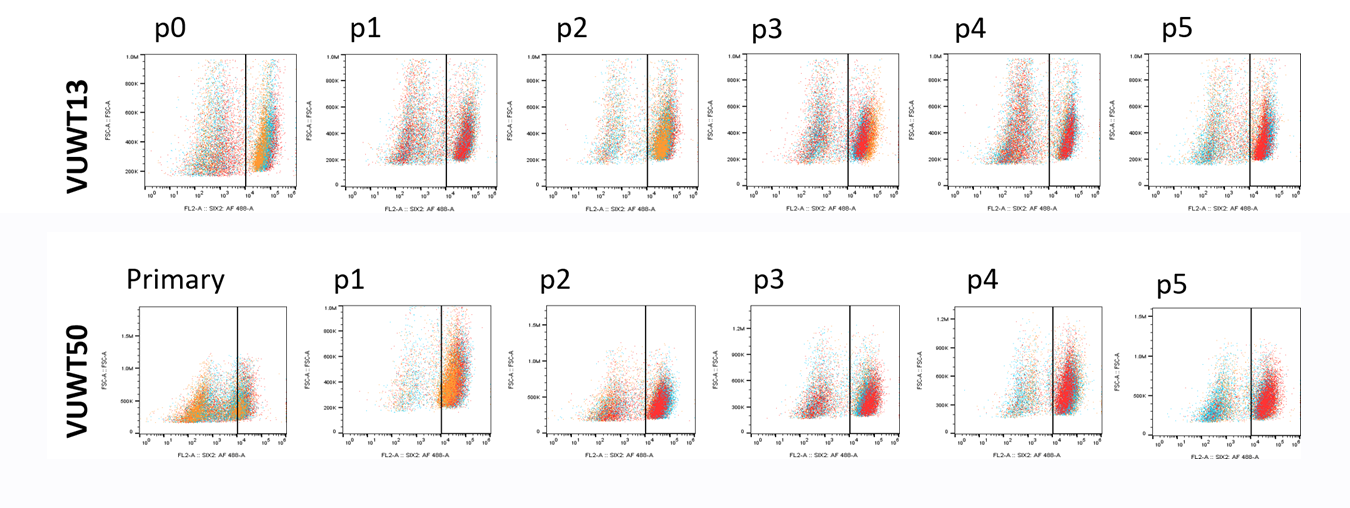

### Supplementary Figure 2

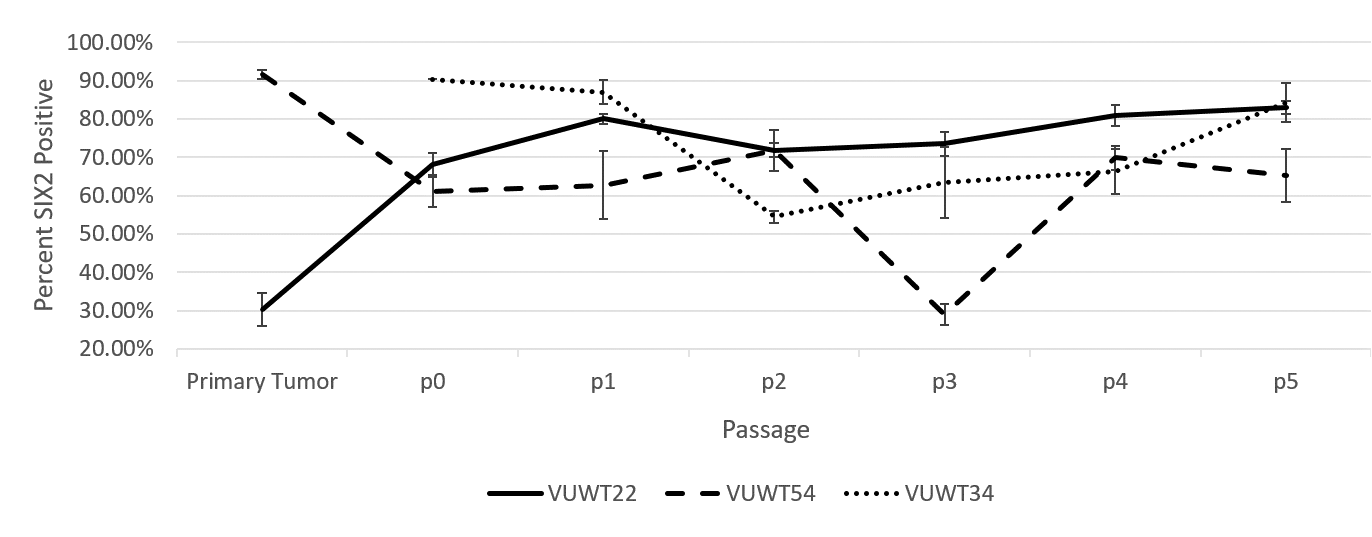

### Supplementary Figure 3

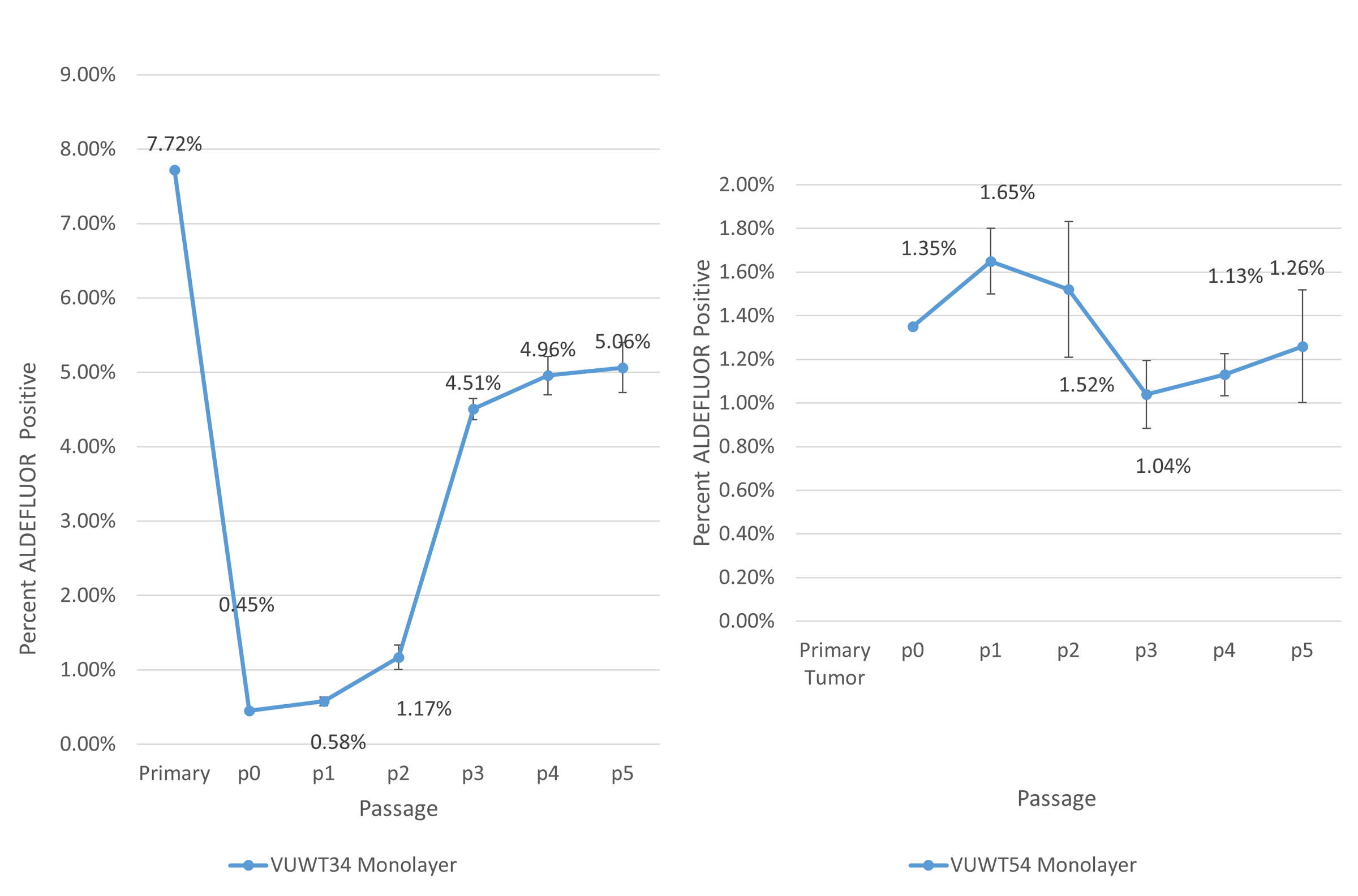

### Supplementary Figure 4

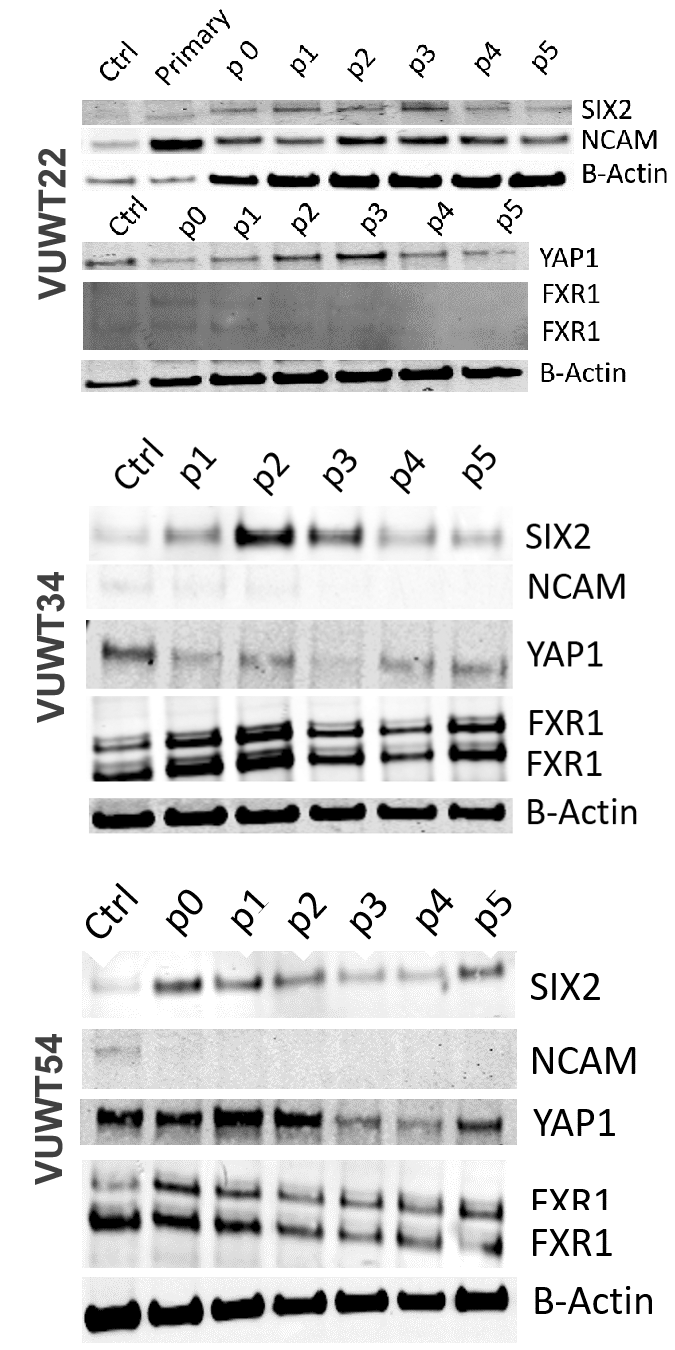

### Supplementary Figure 5A

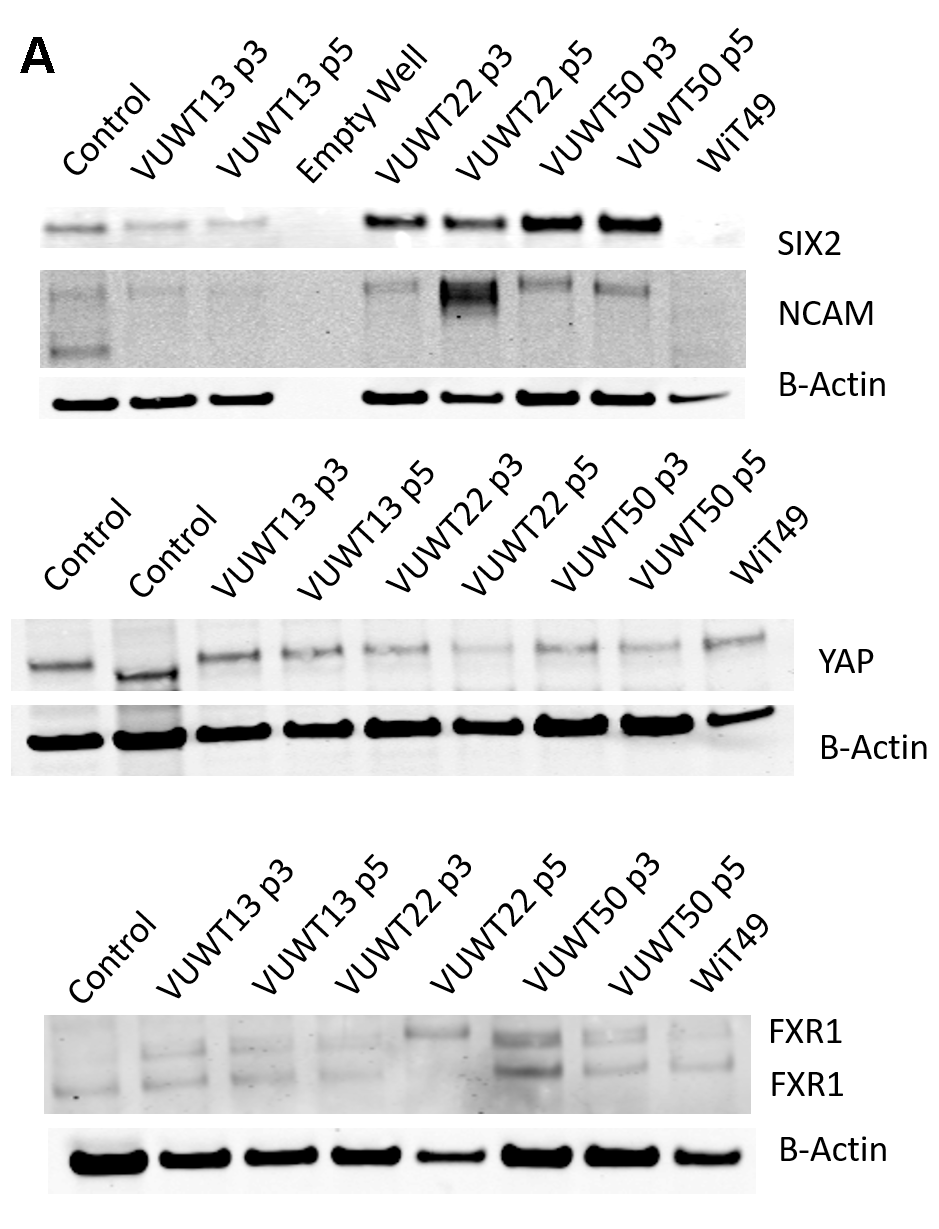

### Supplementary Figure 5BC

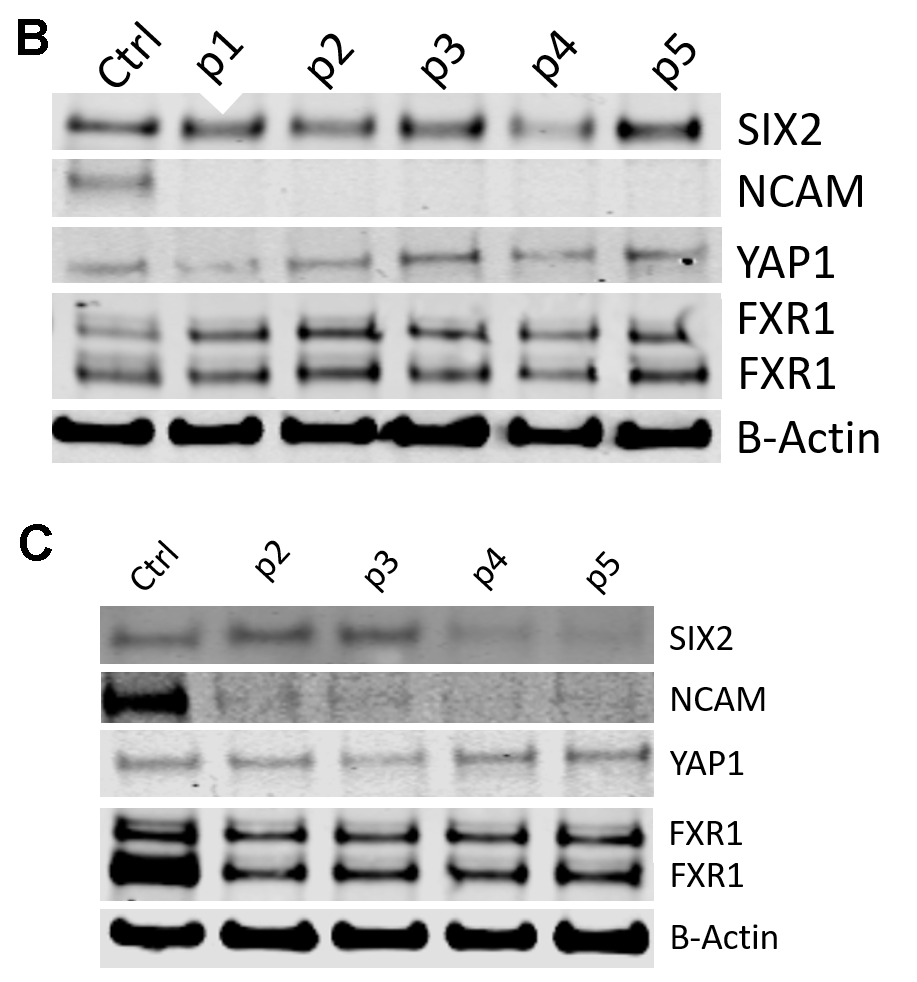
